# Inhibition of HIV-1 gene transcription by KAP1 in myeloid lineage

**DOI:** 10.1101/2020.04.28.066274

**Authors:** Amina Ait Ammar, Maxime Bellefroid, Fadoua Daouad, Valérie Martinelli, Jeanne Van Assche, Clémentine Wallet, Marco De Rovere, Birthe Fahrenkrog, Christian Schwartz, Carine Van Lint, Virginie Gautier, Olivier Rohr

## Abstract

HIV-1 latency generates reservoirs that prevent viral eradication by the current therapies. To find strategies toward an HIV cure, detailed understandings of the molecular mechanisms underlying establishment and persistence of the reservoirs are needed. The cellular transcription factor KAP1 is known as a potent repressor of gene transcription. Here we report that KAP1 represses HIV-1 gene expression in myeloid cells including microglial cells, the major reservoir of the central nervous system. Mechanistically, KAP1 interacts and colocalizes with the viral transactivator Tat to promote its degradation via the proteasome pathway and repress HIV-1 gene expression. In myeloid models of latent HIV-1 infection, the depletion of KAP1 increased viral gene elongation and reactivated HIV-1 expression. Bound to the latent HIV-1 promoter, KAP1 associates and cooperates with CTIP2, a key epigenetic silencer of HIV-1 expression in microglial cells. In addition, Tat and CTIP2 compete for KAP1 binding suggesting a dynamic modulation of the KAP1 cellular partners upon HIV-1 infection. Altogether, our results suggest that KAP1 contributes to the establishment and the persistence of HIV-1 latency in myeloid cells.

## INTRODUCTION

The pandemic of HIV-1 infections is a global health problem. Current cART (combination antiretroviral therapy) efficiently suppresses viral expression below clinical detection levels but fails to eradicate latently infected reservoirs, the main obstacles toward an HIV cure. Indeed, most efforts have focused on understanding latency molecular mechanisms in resting memory CD4+ T-cell reservoirs. However, they are not the only source of viral rebound. Myeloid cells such as monocytes, tissue-resident macrophages and follicular dendritic cells are part of the viral reservoir(1–3). Protected by the blood brain barrier, the central nervous system is a major anatomic reservoir and a sanctuary for the virus (4). Indeed, integrated SIV DNA have been reported in the CNS of infected macaques, with undetectable plasma viral load (5). In the brain, microglial cells are the major reservoirs of latently integrated HIV-1 (6). The specific molecular mechanisms controlling HIV-1 gene silencing in these CNS resident macrophages should be taken into consideration to design strategies toward HIV cure.

We have previously reported the importance of the cellular co-factor CTIP2 (Coup-TF Interacting Protein 2) in the establishment of HIV-1 post-integration latency in microglial cells (7–9). CTIP2 promotes and maintains HIV-1 gene silencing by recruiting chromatin modifying complexes including HDAC1/2 (Histone Deacetylase 1/2), SUV39h1 (Suppressor of Variegation 3-9 Homolog 1) and LSD1 (Lysine-Specific histone Demethylase 1A) to the viral promoter (9, 10). In addition, CTIP2 targets and represses HIV-1 Tat transactivation function by promoting its relocation to heterochromatin structure via the formation of a Tat-CTIP2-HP1α (Heterochromatin Protein 1α) complex (11) and by repressing the elongation factor P-TEFb (12). P-TEFb is the key cofactor of Tat. CTIP2 associates with an inactive form of P-TEFb to repress the CDK9 (Cyclin Dependent Kinase 9) catalytic subunit and inhibit P-TEFb sensitive genes including HIV-1 (12). Finally, we reported that HMGA1 recruits this CTIP2-associated inactive P-TEFb complex to the viral promoter (13). Importantly, Desplats *et al*, revealed that CTIP2 expression is increased in the CSF, astrocytes and microglial cells from PLWH (Patients Living With HIV) on suppressive cART (14). Collectively, our results demonstrate the importance of CTIP2 and its partners in the regulation of HIV-1 gene silencing. The HIV-1 latency is a multifactorial phenomenon involving factors dedicated to epigenetic gene silencing. KAP1 (KRAB (Krugel-Associated Box) domain-Associated Protein 1), also known as TRIM28 (Tripartite Motif-containing Protein 28) or TIF1β (Transcriptional Intermediary Factor 1 beta) is one of them. It was identified in 1996 as an interaction partner of the KRAB-ZFPs (Krüppel-Associated Box Zinc Finger Proteins) transcription factors family members (15). KAP1 is a protein with multiple functional domains that regulates the chromatin environment through interactions with different partners (16). KAP1 is recruited to DNA loci via interactions of its RBCC (RING finger, 2 B-box zinc fingers, Coiled-Coil region) domain with KRAB proteins (17). The RING domain of KAP1 has SUMO and Ubiquitin E3 ligase activities and induces heterochromatin structures (18–20), the PHD (Plant Homeo Domain) C-terminal domain has a SUMO E3 ligase activity needed to SUMOylate its own Bromo domain and the SUMOylated Bromo domain recruits the NURD and SETDB1 repressor complexes (21). KAP1 is known to repress endogenous and latent retrovirus (22). In HIV-1 infected cells, KAP1 associates with HDAC1 to deacetylate the integrase and inhibit proviral integration (23). At the transcriptional level, KAP1 has been reported to contribute to ZBRK1 (Zinc finger and BRCA1 (Breast cancer type 1 susceptibility protein)-interacting protein with A KRAB domain 1) and ZNF10 (Zinc Finger Protein 10)-mediated HIV-1 LTR repression (24, 25). However, these studies have not yet shown the direct involvement of KAP1 and its mechanism of action. Moreover, KAP1-mediated recruitment of an inactive form of P-TEFb to most genes containing a paused RNA polymerase, including the HIV-1 LTR promoter has been described to favor gene expression upon stimulation (26) (27). Despite these controversies, KAP1 is mainly shown as a viral gene repressor. KAP1 restricts the activation of MLV (Murine Leukemia Virus) and HTLV-1 (Human T-Lymphotropic Virus type 1) genes (28, 29). More recently, the restriction factor APOBEC3A (Apolipoprotein B mRNA-Editing enzyme Catalytic polypeptide-like 3G) has been described to recruit KAP1 to suppress HIV-1 transcription (30). In addition, a very recent study suggests that KAP1 represses HIV-1 gene expression by mediating CDK9 SUMOylation resulting in P-TEFb repression in T cells (31). Surprisingly, since KAP1 functions have been extensively studied in T lineage, nothing has been done to define its role and its mechanism of action on HIV-1 infection in myeloid cells.

Here, we studied the influence of KAP1 on HIV-1 expression and its specific mechanism of action in monocytic and microglial cells, the resident macrophages and the main viral reservoirs in the brain. In microglial cells KAP1 repressed HIV-1 gene expression. While KAP1 was found associated with the silenced HIV-1 promoter in the CHME5-HIV microglial model of latency, stimulations released KAP1 from the provirus. RNA interference-mediated knockdown of KAP1 in monocytic cell line latently infected with HIV-1 showed an increase in viral reactivation and transcription. Mechanistically, KAP1 repressed the HIV-1 promoter activity in the absence and in the presence of the viral transactivator Tat. Moreover, KAP1 had a major repressive impact on Tat function. We report that KAP1 interacts physically and colocalizes with Tat in the nucleus of microglial cells to promote its degradation via the proteasome pathway. Finally, KAP1 was found associated with the repressor CTIP2. They both cooperated to repress Tat function. Altogether our results highlight the contribution and the specific mechanism of action of KAP1 in the establishment and the persistence of HIV-1 post-integration latency in cells from myeloid origin.

## MATERIAL AND METHODS

### Plasmids

The following plasmids: pcDNA3, flag CTIP2, flag-CTIP2 350-813, 350-717, 1-354 and 145-434, Tap-CTIP2, flag-Tat, DsRed-Tat, ShCTIP2, LTR-Luc, pNL4-3Δ Env Luc have been described previously (9, 11, 12). The flag-KAP1 plasmids, flag-KAP1 S824A/D come from Dr. David K. Ann (32). The vectors shKAP1 and shNT (non-target) were gift from Dr. Ivan D’orso (26). The plasmids flag-KAP1WT, flag-KAP1 ΔRBCC, ΔPHD, ΔBromo, ΔPHD/ΔBromo were provided by Dr. Florence Cammas (IRCM Montpellier). GFP-KAP1 comes from Addgene (#65397).

### Cell culture

The human microglial cell line (provided by Prof. M. Tardieu, Paris, France) (33), HEK293T cell lines and CHME5 cell lines latently infected with HIV-1 (34) were maintained in Dulbecco’s modifed Eagle medium (DMEM) containing 10% decomplemented fetal calf serum and 100 U/ml penicillinstreptomycin. When indicated, the cells were treated with 50 μM MG132 for 6 hours. THP89 cell lines were grown in RPMI 1640 medium supplemented with 10% decomplemented fetal calf serum and 50 U/ml penicillin-streptomycin.

### Antibodies and reagents

The list of antibodies and other chemical reagents used in this work are listed in the following table 1.

| Reagent | Reference |
| --- | --- |
| anti-CTIP2 antibody | A300-383A Bethyl |
| anti-KAP1 antibody | Ab10483 Abcam |
| anti-Tat antibody | ab43014 Abcam |
| anti-β-actin antibody | A1978 Sigma-Aldrich |
| anti-Flag antibody | F3165 Sigma-Aldrich |
| anti-GFP antibody | 632592 Clontech |
| anti-α-tubulin antibody | Ab4074 Abcam |
| anti-UBC9 antibody | 610749 BD Biosciences |
| anti-HA antibody | 901 501 Biolegend |
| anti-mouse IgG-HRP | Sc-358914 Santa Cruz |
| anti-rabbit IgG-HRP | Sc-2004 Santa Cruz |
| anti-KAP1 ChIP grade antibody | C15410236-100 Diagenode |
| anti-RNA Pol II ChIP grade antibody | 14958 Cell signaling |
| Rabbit IgG | C15410206 Diagenode |
| Mouse IgG | C15400001-15 Diagenode |
| Reverse PCR | 1708890 Bio Rad |
| Quantitative PCR | Ssoadvanced™ Universal SYBR Green Supermix Bio Rad |
| anti-proteases | 000000011873580001 Sigma-Aldrich |
| MG132 | BML-PI102-0025 ENZO Life Sciences |
| TNFα | 300-01A PeproTech |
| HMBA | 224235 Sigma-Aldrich |
| DMA | 20660 Thermoscientific |
| anti-flag immunoprecipitation beads | A-2220 Sigma-Aldrich |
| Protein G magnetic beads | C03010021-220 Diagenode |
| Protein A magnetic beads | C03010020-220 Diagenode |
| Protein A Agarose/Salmon Sperm DNA | 16-157 Millipore |

### Lentiviral production

HEK cells seeded at a density of 4X10^6^ and grown in a 10 cm dish were transfected with the different shRNAs (shNT or shKAP1)-containing plasmids (9 μg), the pVSV-G (2.25 μg) and the psPAX2 (6.75 μg) packaging vectors by the calcium phosphate transfection method according to the manufacturer’s protocol (TaKaRa; Calphos). 72h post-transfection, the virus-containing supernatant (8ml) was filtered (Steriflip; Merck) before being concentrated 20x (14000 rpm, 1h30, 4°c) and stored at −80°c.

### Generation of stable THP89 knock-down for KAP1

THP89 cells were transduced by spinoculation as previously described (26). Briefly, 100μl of 20X concentrated lentiviral stock was added to 10^6^ cells previously washed and resuspended in 400μl of complete growth medium containing polybrene (8μg/ml). Cells were then centrifuged (3200 rpm 1h, 32°c) before being incubated for 2h at 37°c. After this incubation, cells were washed and transferred in 1 ml complete growth medium in a 12-well dish. 48h post-infection, infected cells were selected by puromycin selection at a final concentration of 1 ug/ml for 2 weeks. GFP positive cells were measured by Flow Cytometry. Total proteins and RNAs were then extracted as described above for RT-qPCR and western blotting.

### Nuclear protein extract

Cells seeded at a density of 3×10^6^ and grown in a 10 cm dish were transfected with the indicated vectors (30μg) using the calcium phosphate coprecipitation method. 48 hours post-transfection, cells were lysed for 10 minutes on ice in a buffer containing: 10 mM HEPES, 1.5 mM MgCl2, 10 mM KCl and 0.5 mM DTT. After one minute of centrifugation at 13000 g, the nuclear pellet was lysed for 30 minutes on ice in another buffer containing: 20 mM HEPES, 25% glycerol, 1.5 mM MgCl2, 420 mM NaCl, 0.2 mM EDTA, 0.5 mM DTT. After 2 minutes of centrifugation at 13000 g, the supernatant was analyzed by immunoprecipitation and Western blotting. Anti-proteases were systematically added to the lysis buffers.

### Total protein extract

Cells seeded at a density of 10^6^ cells per well and grown in a 6-well dish were transfected using the calcium phosphate coprecipitation method with the indicated vectors (12 μg). After 24 hours, the cells were lysed on ice for 50 minutes using a buffer containing: 10 mM Tris HCl pH 7.5, 150 mM NaCl, 1 mM EDTA, 1% NP40, 0.5% Na deoxycholate, 0.1% SDS. After 10 minutes centrifugation at 13000 g, the supernatant was stored for Western blot analysis. Antiproteases were systematically added to the lysis buffers.

### Co-immunoprecipitation and Western blot analysis

Immunoprecipitations were performed on 500 μg of nuclear protein extracts using anti-flag M2 gel (Sigma-Aldrich) or by using Diagenode A/G protein-coupled magnetic beads to target endogenous proteins. Briefly, the nuclear protein extract was cleared for 1 hour under rotation at 4°C with the A/G protein-coupled magnetic beads. Meanwhile, the beads were coated under rotation with 1 μg of antibody for 30 minutes at room temperature. After the preclearing step, the beads were removed, replaced by the antibody coated beads and incubated overnight at 4°C under rotation. The day after, the incubated beads were washed 2 times with a low salt buffer (50mM TrisHCl pH 7.5, 120 mM NaCl, 0.5mMEDTA, 0.5% NP40, 10% glycerol), 2 times with a high salt buffer (50mM TrisHCl pH 7.5, 0.5M NaCl, 0.5mM EDTA, 0.5% NP40, 10% glycerol) and one time with a low salt buffer. The immunoprecipitated proteins were eluted by heating at 100°C during 15 minutes in 4x Laemmli Sample Buffer (BIORAD). Immunoprecipitated protein complexes were subjected to Western blot analysis. The proteins were detected using the antibodies as indicated and visualized by a Thermo Scientific Chemiluminescence Detection System (Pierce ™ ECL Western Blotting Substrate 32106).

### Fluorescence Microscopy

The microglial cells were seeded at 0.8×10^6^ and grown on coverslip in a 24-well plate. The cells were transfected after 24 hours with the indicated vectors by the Jetprime (Polyplus transfection) method, according to the instructions of the supplier. After 48 hours of transfection, the cells were fixed with PBS containing 2% formaldehyde for 15 minutes. The cells were then washed 3 times with PBS. The coverslip was mounted on a slide using the Mowiol 4-88 (Sigma-Aldrich) containing 1μg/ml DAPI. Cells fluorescence was observed under a fluorescence microscope and imaged using a Zeiss Axio Observer inverted microscope (Zeiss, Oberkochen, Germany) with Zeiss Plan-Apochromat 100x/1.4 oil objective. Images were acquired using the microscope system software AxioVision V 4.8.2.0 and processed using Image J software.

### Luciferase assay

Microglial cells, seeded at 0.4×10^6^ in a 48-well dish, were transfected in triplicate by the calcium phosphate coprecipitation method with the indicated vectors. 48 hours post-transfection, the cells were collected and the luciferase activity was measured using the Dual-Glo Luciferase Assay system (Promega Madisson, USA) and normalized to Renilla Luciferase activity.

### Chromatin immunoprecipitation assay

Chromatin immunoprecipitation assay was performed following ChIP assay kit (EMD Millipore) protocol. Briefly, HIV-1 CHME5 cells were seeded at 3×10^6^ in T175 Corning flask. 24 hours later, the cells were mock treated or stimulated with 10 ng/ml of TNFα (Tumor Necrosis Factor α) and 5 mM of HMBA (Hexamethylene bisacetamide). After 24 hours the cells were trypsinated, counted and then washed with PBS. The cells were resuspended at ratio of 4×10^6^ cell/ml of PBS. The cells were cross-linked with a freshly made DMA (dimethyl adipimidate) at 5 mM for 25 minutes followed by a second 8 minutes cross-link with formaldehyde at 1%. The cross-linking reaction was stopped by 0.125 M of Tris-Glycine solution. Cells were washed twice with ice cold PBS then resuspended to a ratio of 10×10^6^/300μl of SDS Lysis buffer (Tris HCl 50 mM pH 8, EDTA 10 mM, 1% SDS) containing proteases inhibitors. The chromatin was sonicated (Bioruptor Plus, Diagenode) 40 minutes (30s on, 30s off) to obtain DNA fragments of 200–400 bp. Chromatin immunoprecipitations were performed with chromatin from 5×10^6^ cells and 5 μg of antibodies. IgG was used as a control for immunoprecipitation. The DNA was purified using the High Pure PCR Product Purification Kit (11732676001 Sigma-Aldrich). Quantitative Real Time PCR reactions were performed with the TB Green™ Premix Ex Taq™ II (TAKARA) as recommended by the supplier. Relative quantification using standard curve method was performed for the primer pair, and 96-well plates (MU38900 Westburg) were read in a StepOnePlus PCR instrument (Applied Biosystems). The results are represented as a fold enrichment relative to the IgG condition. The LTR5’ forward and reverse Primer sequences used are respectively: 5’-GCTTAAGCCTCAATAAAGCTTGC-3’, 5’-TGACTAAAAGGGTCTGAGGGAT-3’.

### Flow cytometry

CHME5 HIV-1 cells mock treated or stimulated with 10 ng/ml of TNFα and 5 mM of HMBA for 24 hours or either THP89 HIV-1 cells transduced with lentiviral particles were washed twice in PBS, resuspended in PBS containing 4% paraformaldehyde and fixed for 1 hour at 4°C in the dark. Cells were then washed twice in PBS and resuspended in FACS buffer (PBS, 0.1% BSA, 0.1% NaN3). The percentage of GFP-positive cells was measured on a CXP cytometer (Cytomics FC 500, Beckman Coulter) using CXP Software version 1.0 according to the manufacturer’s instructions.

### mRNA quantification

Total RNA was extracted from cells expressing the indicated vectors according to the QIAGEN RNeasy Plus Kit protocol or with Tri-Reagent (TRC118, MRC) according to the manufacturer’s instructions. Following DNAse treatment (AM1907, Invitrogen), the cDNAs were produced using BioRad Reverse Transcription Kit (iScriptTM Reverse Transcription Supermix) or the PrimeScript RT reagent kit (RR037A, TaKaRa). cDNAs from HEK and THP89 cells were quantified by quantitative PCR using Bio Rad’s SsoAdvanced Universal SYBR Green Supermix. Data were calculated using the 2^-(ΔΔCT)^ method and normalized to the amount of GAPDH.

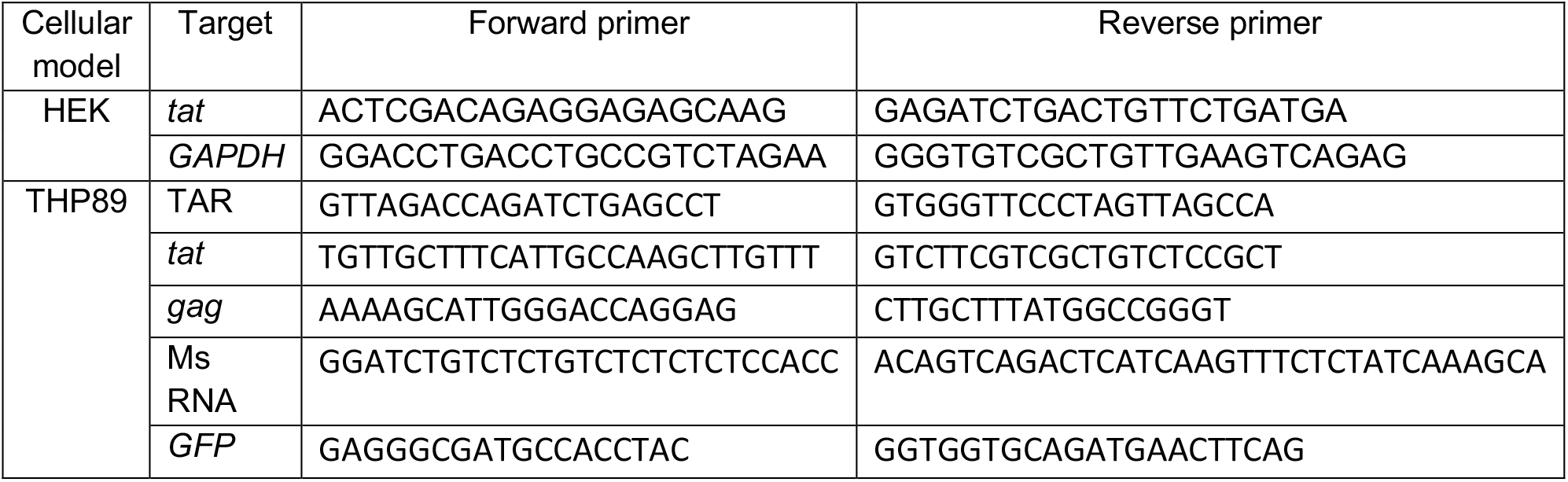

### Statistical analysis

Data were analyzed by performing the student t-test using Microsoft Excel. Values of *p* <0.05 (*), *p* <0.01 (**) and *p* <0.001 (***) were considered significant. The results were represented as mean and standard deviation of at least 3 independent experiments.

## RESULTS

### KAP1 cooperates with CTIP2 to repress HIV-1 gene expression in microglial cells

The functional impact of KAP1 overexpression or depletion on HIV-1 gene expression was examined on microglial cells expressing the pNL4-3 Δ ENV Luc HIV molecular clone (Figure 1 A and 1 B) or the episomal vector LTR-Luc (Figure 1 C and 1 D). In both cases we observed that expression of increasing amounts of KAP1 inhibited the HIV-1 promoter activity in a dose-dependent manner, while KAP1 depletion stimulated its activity.

**Figure 1:**
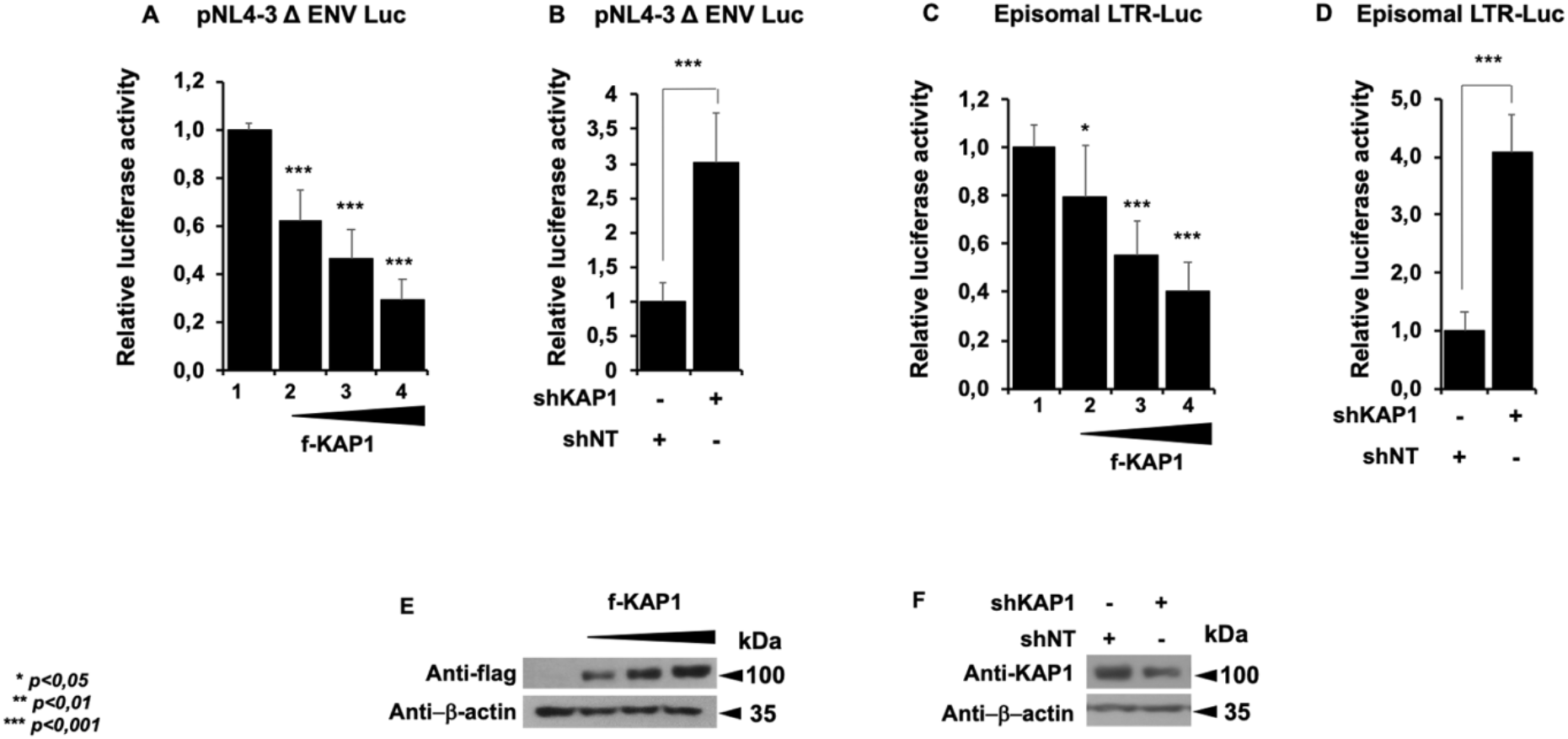
KAP1 suppresses HIV-1 expression. (A-D) Microglial cells were transfected with the pNL4-3 Δ ENV Luc vectors (A, B) or with episomal vector LTR-Luc (C, D) in the presence of an increasing dose of KAP1 (A, C), or with a KAP1 knockdown (shKAP1) (B, D). Luciferase activity was measured 48 hours post-transfection and after lysis of the cells. Luciferase values were normalized to those obtained with LTR-Luc or pNL4-3 Δ ENV Luc alone. A t-test was performed on 3 independent experiments *P* (* *p* <0.05, ** *p* <0.01, *** *p* <0.001). (E, F) Overexpressions of KAP1, as well as knockdown efficiencies, were validated by Western blotting.

These results are consistent with a transcriptional control of the viral expression by KAP1 and support a repressive role of KAP1 on HIV-1 gene expression. Of note, KAP1 protein expression levels were assessed by Western blot (Figures 1 E and F). Given KAP1 capacity to silence HIV-1 gene expression, we next examined the presence of KAP1 at the silenced viral promoter in CHME5-HIV microglial model of HIV-1 latency (34). Chromatin IP experiments revealed the presence of KAP1 associated with the silent 5’LTR region of the provirus (Figure 2 A). However, treatments with TNFα and HMBA to induce viral reactivation significantly displaced KAP1 from the promoter (Figure 2 A). The activation of HIV-1 promoter was validated by increased binding of the RNA Pol II (Figure 2 A) and by quantification of the GFP positive cells (Figure 2 B). Our results suggest that KAP1 targets the latent viral promoter to promote HIV-1 post-integration latency in microglial cells. To confirm these results in other models of infected myeloid cells, we took advantage of THP89 cells latently infected with recombinant p89.6 HIV-1 reporter virus wherein the GFP gene is inserted in the viral genome(35). We generated THP89 monocytic cell lines expressing nontarget (shNT) or KAP1 (shKAP1) small hairpin (sh) RNAs. The efficiency of KAP1 knockdown in the selected puromycin resistant cells was validated by Western blot (Figure 3 A). Using these cells, we assessed the role of KAP1 in viral latency and reactivation by monitoring GFP expression. Flow cytometry analysis and GFP transcripts quantifications revealed an increase of GFP positive cells upon KAP1 knockdown (Figure 3 A and B). Activation of HIV-1 gene transcription in THP89 cells following KAP1 depletion was further studied by quantifying initiated transcripts (TAR region), elongated transcripts (*gag* and *tat* regions), and the multiply spliced transcripts (Ms RNA) (Figure 3 C). Significant higher levels of elongated and multiply-spliced transcripts were observed in KAP1 knocked-down cells. Altogether, our results indicate that KAP1 depletion enables the release of HIV-1 transcriptional blocks underlying latency.

**Figure 2:**
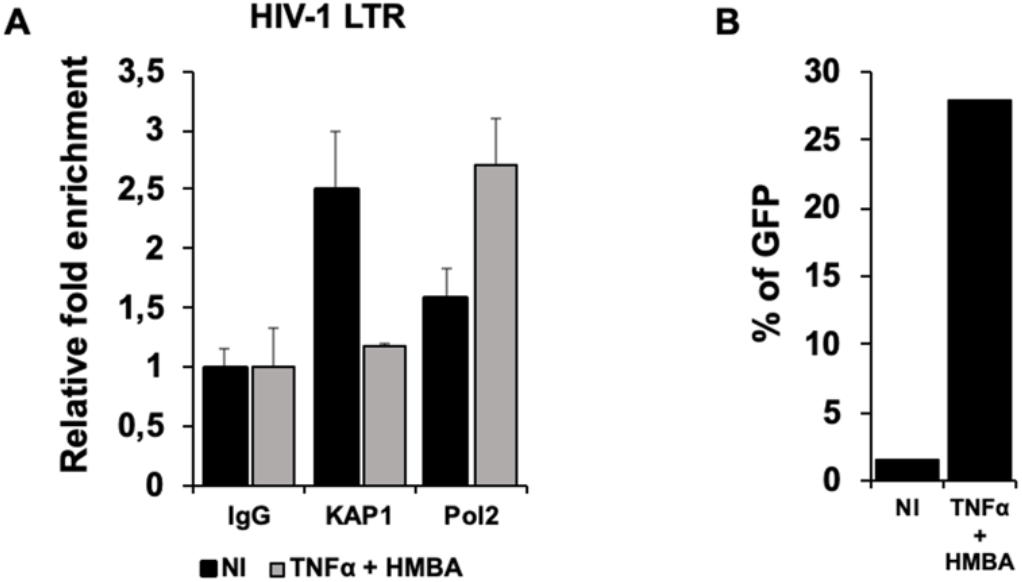
KAP1 binds to the latent HIV-1 promoter. (A) Microglial cells latently infected with HIV-1 containing the GFP reporter gene were treated for 24h with TNFα and HMBA. Cells were subjected to ChIP-qPCR experiments targeting the binding to the HIV-1 5’ LTR promoter (B) The same microglial cells were also analyzed by flow cytometry. This figure is representative of 3 independent experiments.

**Figure 3:**
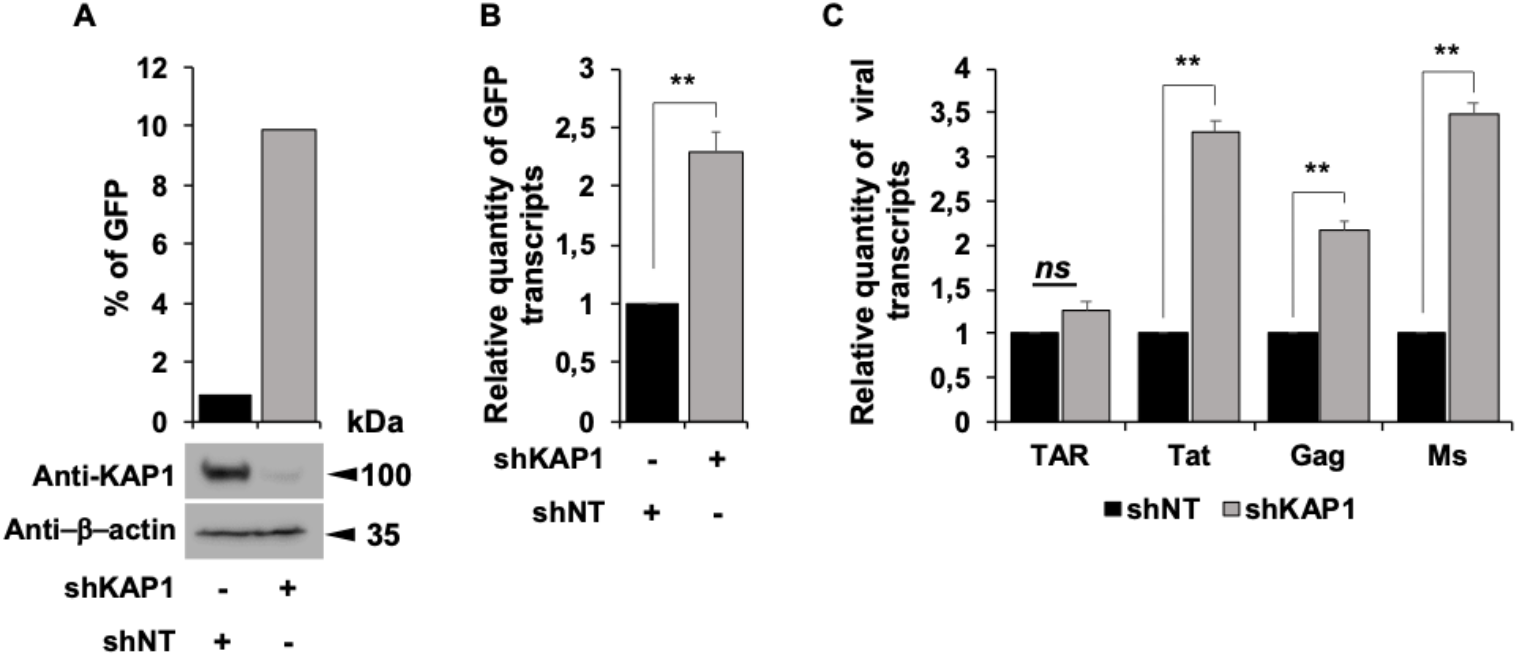
Reactivation of HIV-1 gene transcription in myeloid models of latency upon KAP1 depletion. (A-C) The HIV-1 infected monocytic (THP89) cells containing a GFP reporter gene were transduced either with non-targeting shRNA (indicated as shNT) or shKAP1. (A) The GFP expression was examined by flow cytometry and the KAP1 protein expression level was assessed on total cellular extract by western blot. (B, C) Total RNA products from shNT or shKAP1 THP89 transduced cells were retrotranscribed. GFP and HIV-1 gene transcripts as Initiated (TAR region), elongated (*gag*, *tat*), and multiple-spliced RNA (Ms RNA) were quantitated by real–time RT-PCR. The relative mRNA level was first normalized to GAPDH then to the shNT-transduced condition. Results are means from duplicate.

We have previously reported the importance of the cellular cofactor CTIP2 in the establishment and the persistence of HIV-1 latency (9, 10, 13). Since KAP1 and CTIP2 have been both involved in the control of HIV-1 gene expression, we next investigated whether they interact physically and functionally. Looking at the viral expression level, we observed that the concomitant overexpression of KAP1 and CTIP2 had a significantly stronger repressive activity than when overexpressed individually, (Figure 4 A, column 4 versus 2 and 3). Similarly, we observed a modest but significant cooperation in the context of a simultaneous depletion of the two proteins (Figure 4 B, column 4 versus 2 and 3). As shown in figure 4 C and 3 D, similar results were obtained upon analysis of the activity of 5’LTR viral promoter. To ensure that the effects can be correlated to the expression levels of CTIP2 and KAP1, we performed Western blot analysis. The results demonstrated the efficiencies of the overexpression and the knockdown at the protein level (Figures 4 E and F). We have reported that CTIP2 represses Tat function by relocating the viral regulatory protein in heterochromatin structures (11). We therefore investigated the impact of KAP1 on Tat function. As expected, Tat expression stimulated the viral promoter activity (Figure 5 A, column 2). However, while the overexpression of KAP1 repressed this transactivation (Figure 5 A, columns 3 to 5 versus 2), the knockdown of KAP1 promoted Tat activity (Figure 5 B). These results suggested us that the repressive function of KAP1 on HIV-1 expression results from a double impact on the initiation (-Tat) and the elongation (+Tat) steps of viral gene transcription. We next modulated the expression of KAP1 together with CTIP2. As expected, overexpression of KAP1 and CTIP2 alone inhibited Tat function (Figure 6 A, column 3 and 4). Moreover, their concomitant overexpression further repressed Tat-mediated activity (figure 6 B, column 4). As shown in Figure 6 C and 6 D, the efficiencies of the overexpression and the depletions were controlled by Western blot. Taken together, our results demonstrate that KAP1 and CTIP2 cooperate functionally to repress HIV-1 gene expression in microglial cells.

**Figure 4:**
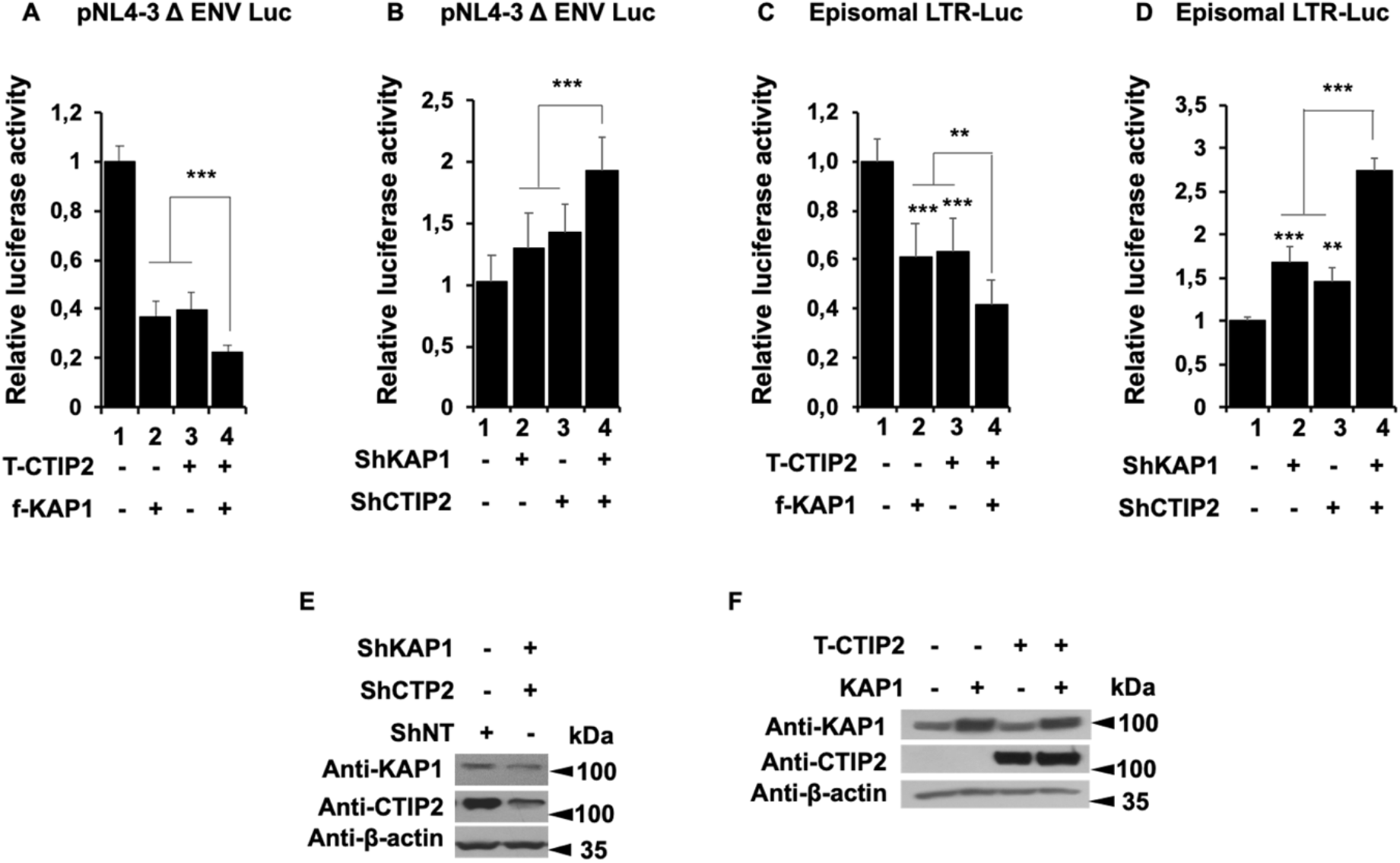
Kap1 and CTIP2 together contribute to the silencing of HIV-1 gene expression. (A-D) Microglial cells were transfected with the pNL4-3 Δ ENV Luc vectors (A, B) or LTR-Luc (C, D) under the indicated conditions. The cells were lysed after 48h of transfection. The luciferase activities were measured and normalized to the conditions where the pNL4-3 Δ ENV Luc and LTR-Luc vectors were transfected alone. A t-test was performed on 3 independent experiments *P* (* *p* <0.05, ** *p* <0.01, *** *p* <0.001). (E, F) Over-expression and knockdown of the indicated proteins were validated by Western blot.

**Figure 5:**
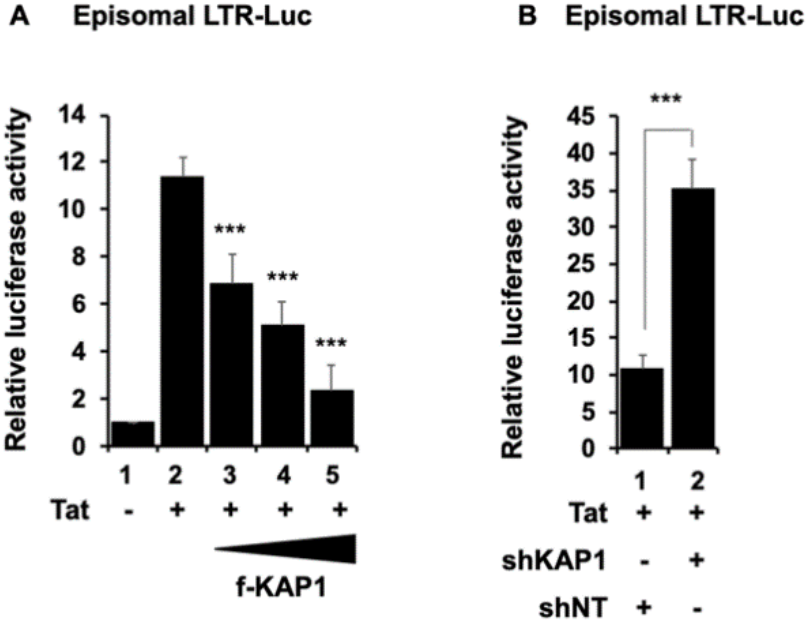
KAP1 inhibits Tat activity. (A, B) Microglial cells were transfected with LTR-Luc vector under the indicated conditions. After 48h of transfection, the luciferase activities were measured after cell lysis and normalized under conditions where LTR-Luc vector were transfected alone. A t-test was performed on 3 independent experiments *P* (* *p* <0.05, ** *p* <0.01, *** *p* <0.001).

**Figure 6:**
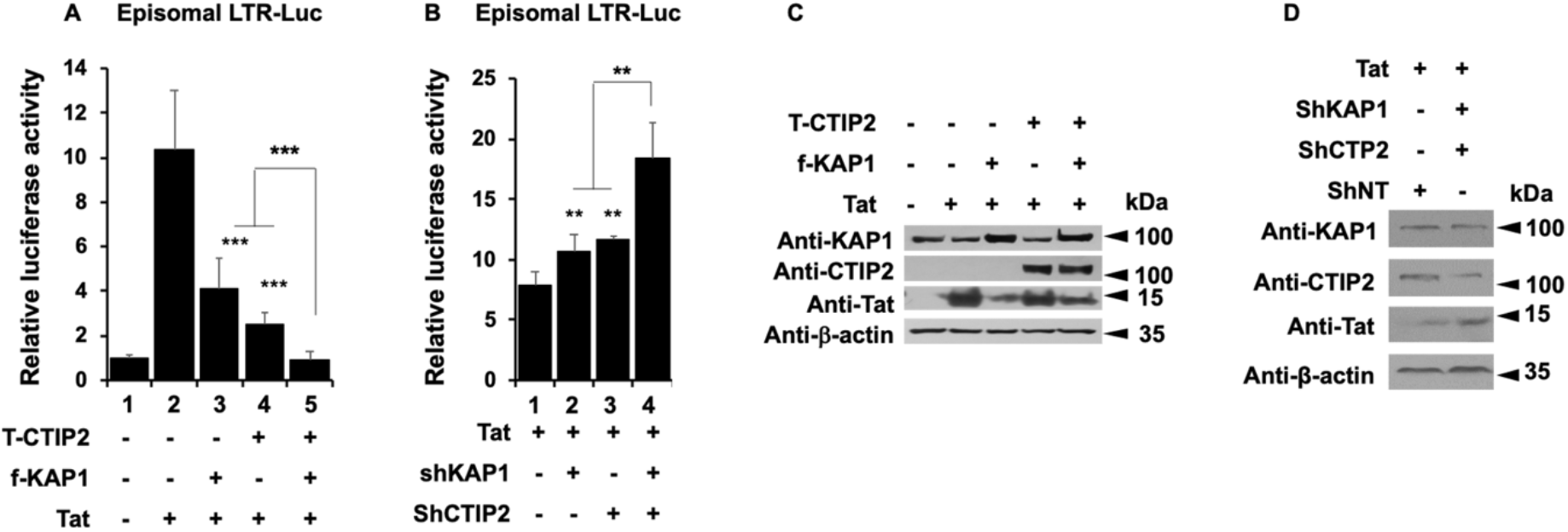
KAP1 cooperates with CTIP2 in Tat repression. (A, B) Microglial cells were transfected with LTR-Luc vector under the indicated conditions. After 48h of transfection, the luciferase activities were measured after cell lysis and normalized under conditions where LTR-Luc vector was transfected alone. A t-test was performed on 3 independent experiments *P* (* *p* <0.05, ** *p* <0.01, *** *p* <0.001). (C, D) Over-expression and knockdown of the indicated proteins were validated by Western blot.

### KAP1 interacts and colocalizes with Tat

Since our results suggest a physical interaction between Tat and KAP1, we performed coimmunoprecipitation targeting ectopically expressed KAP1 (Figure 7 A) and endogenously expressed KAP1 (Figure 7 B). We show that Tat associated with KAP1 in both conditions. To next visualize the association of KAP1 and Tat in the nucleus, microglial cells expressing GFP-KAP1 and DsRed-Tat were analyzed by direct fluorescence microscopy. As previously described in the literature, KAP1 was found in the nucleoplasm and excluded from the nucleoli, (Figure 7 C upper panel) (36). Tat localized in both nucleoplasm and nucleoli, (Figure 7 C, middle panel) (37, 38). We observed that Tat colocalizes with KAP1 in the nucleoplasm but not in the nucleoli, (Figure 7 C, lower panel).

**Figure 7:**
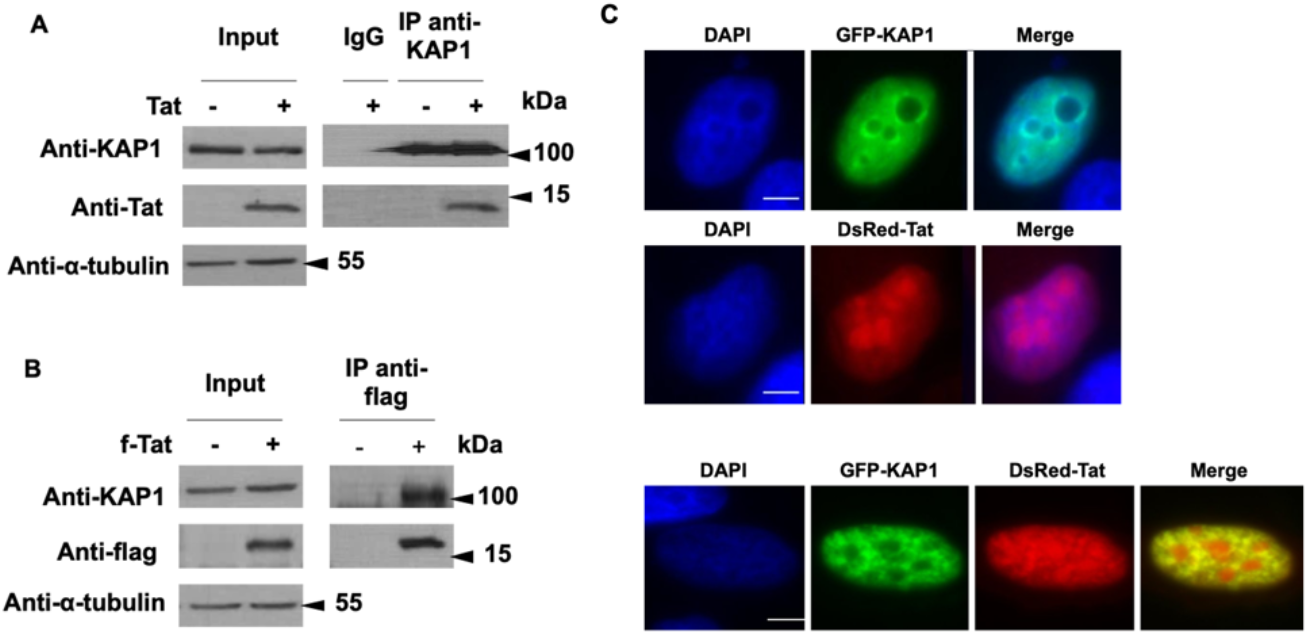
KAP1 interacts and colocalize with Tat. (A, B) HEK cells were transfected with the indicated vectors. After 48h of transfection, the nuclear protein extracts were immunoprecipitated with KAP1 (A) and flag (B) antibodies. The eluted protein complexes were analyzed by Western blotting for the presence of the indicated proteins. (C) Microglial cells were transfected for 48h with the indicated vectors and subjected to observation under a fluorescence microscope. Scale bars are set to 5μm.

### KAP1 promotes Tat degradation by the proteasome pathway

To further characterize mechanistically the repression of Tat by KAP1, we observed the pattern of Tat expression in the presence of increasing amount of KAP1 (Figure 8 A). We found that Tat expression was inversely correlated to the protein level of KAP1 upon KAP1 overexpression (Figure 8 A) and KAP1 depletion (Figure 8 B). To determine if these modulations of Tat protein expression are controlled post-transcriptionally, we quantified Tat mRNAs by RT-qPCR, in the presence or lack of KAP1 overexpression. Since the amount of Tat mRNA remained unaffected by KAP1 (Figure 8 C), we hypothesized that KAP1 overexpression leads to Tat depletion by regulating the stability of the protein. Among the mechanisms controlling protein half-life in eukaryotic cells, the degradation via the proteasome pathway has been long studied. Interestingly, Tat has been described to be directed towards proteasome degradation (39–44). To determine whether the reduced expression of Tat in the presence of KAP1 is related to proteasome degradation, we quantified Tat expression levels in the presence of KAP1 and the proteasome inhibitor MG132, (Figure 8 D). As expected, in the absence of MG132, KAP1 expression reduced significantly Tat expression (Figure 8 D, column 2 versus 1). However, MG132 treatment increased Tat protein levels by about 50% (Figure 8 D column 3 versus 1) and counteracted KAP1-mediated degradation of Tat (Figure 8 D, column 4 versus 2). Thus, our results suggest that KAP1 targets Tat to degradation via the proteasome pathway.

**Figure 8:**
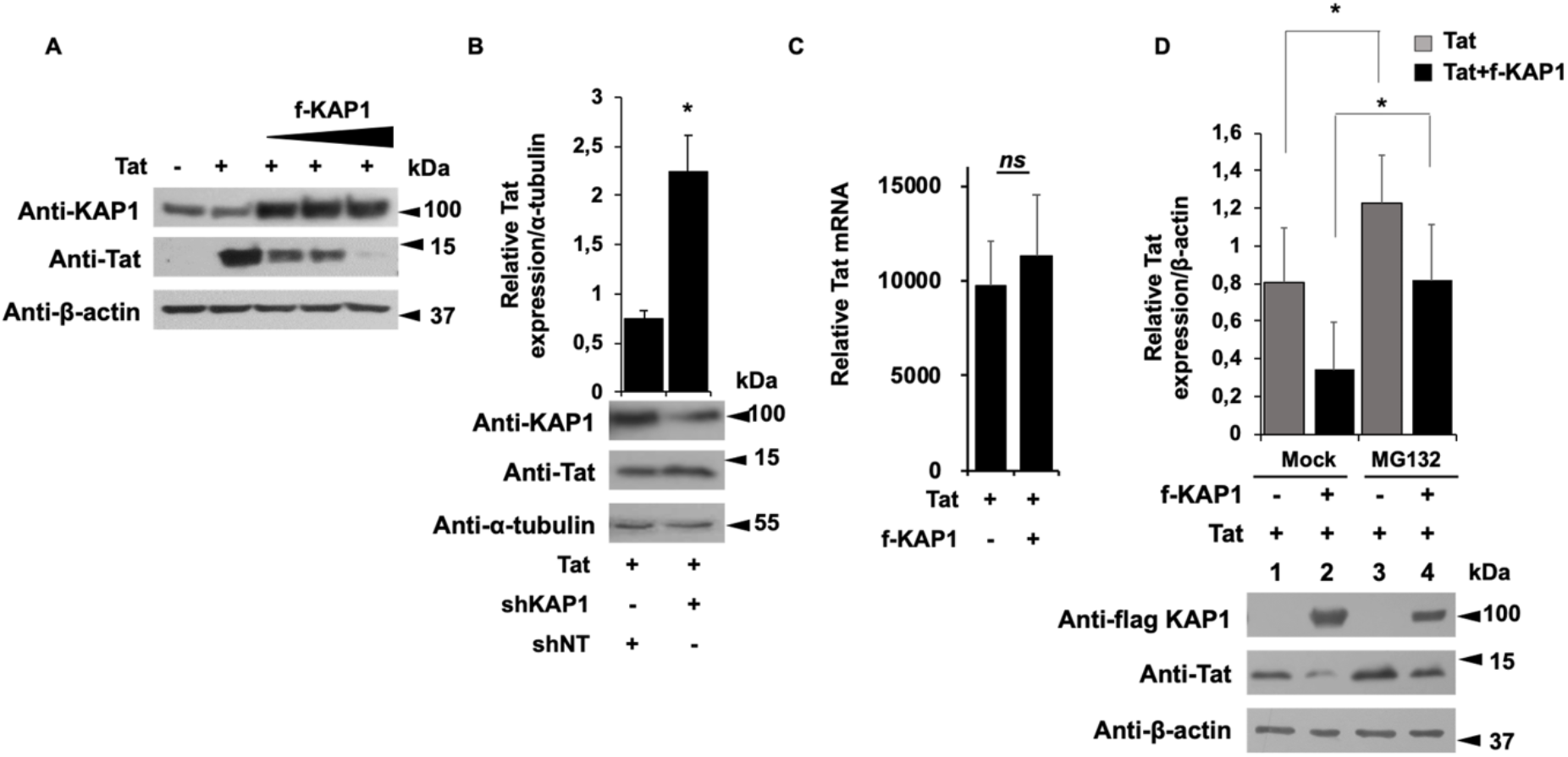
KAP1 promotes Tat degradation via the proteasome pathway. (A, B) HEK cells were transfected with the indicated vectors. 48h later, the nuclear proteins were analyzed by Western blot for the presence of KAP1 and Tat. (B) Tat expression level whose relative quantification to the α-tubulin was carried out using the image J software. (C) The RNA extracts from HEK cells transfected with the indicated plasmids were submitted to RT-qPCR experiments against Tat. The relative mRNA level of Tat was normalized to the GAPDH gene. (D) HEK cells were transfected with the indicated vectors. After 18h of transfection, the cells were treated or not with 50 μM of MG132 for 6h. 24h post-transfection, the total protein extracts were analyzed by Western blot for the presence of KAP1 and Tat. Tat expression level was quantified relatively to the β-actin expression, using image J software. A t-test was performed on 3 (B, C) and 5 (D) independent experiments *P* (* *p* <0.05, ** *p* <0.01, *** *p* <0.001).

### Tat degradation is mediated by the PHD and Bromo domains of KAP1

Using different KAP1 deletion mutants, we next identified the KAP1 domain required for Tat degradation (Figure 9 A). While deletion of the RBCC domain (ΔRBCC) did not affect KAP1-mediated degradation of Tat (Figure 9 A, columns 2 and 3), deletions of the PHD (ΔPHD) and the Bromo domain (Δ Bromo) impaired Tat degradation (Figure 9 A, columns 4, 5 and 6 versus 2). We next used two mutants of KAP1, one mimicking a constitutive phosphorylation of Serine 824 (S824D) and the second which results in a non-phosphorylated form of AKP1 (S824A), (Figure 9 B). Of note, the phosphorylated status of the Serine 824 determine the repressive activity of KAP1 and its ability to interact with the NuRD and SETDB1 repressor complexes via the Bromo domain (32). While expression of KAP1 S824A promoted the degradation of Tat (Figure 9 B, column 5 versus 2), the S824D mutant did not (Figure 9 B, column 4). Overall, these results suggest that KAP1-mediated degradation of Tat may be controlled by phosphorylation of the protein and thereby may depend on the global physiology of the infected cells.

**Figure 9:**
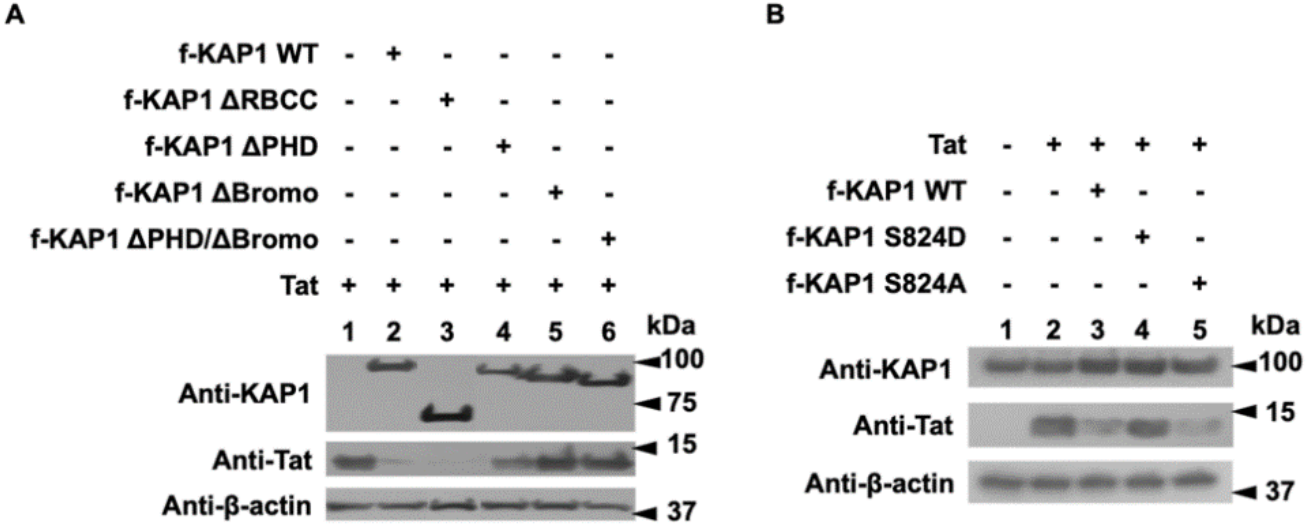
Tat degradation is mediated by the PHD and Bromo domains of KAP1. (A, B) HEK cells were transfected as indicated. 48h later, the nuclear proteins were extracted and tested by Western blot for the expression of the indicated proteins.

### CTIP2 is interacting with KAP1 and competes with Tat for KAP1 binding

We demonstrated that KAP1 functionally cooperates with CTIP2 to inhibit HIV-1 gene expression. Since both CTIP2 and KAP1 have been reported to be associated with an inactive form of P-TEFb, we hypothesized that their functional interactions may result from physical association (12, 26). Immunoprecipitations targeting CTIP2 demonstrate that KAP1 associates with CTIP2 in RNA independent manner (Figure 10 A, column 4 and 6). Similar results were obtained after KAP1 immunoprecipitation (Figure 10 B, column 8 and 10) in the context of ectopic protein expression. We next confirmed these observations by immunoprecipitations targeting the endogenous proteins (Figure 10 C). To delineate the interface between KAP1 and CTIP2, we used different truncations of KAP1 and CTIP2 (Figure 10 D and E). We found that CTIP2 interacts with ΔPHD, ΔBromo, ΔPHD/ΔBromo deleted KAP1 (Figure 10 D, columns 10, 11, 12), but not with the ΔRBCC KAP1, (Figure 10 D, column 9). On the other hand, KAP1 binds only the 1-354 and 145-434 truncated forms of CTIP2 (Figure 10 E, column 11 and 12). Altogether, we found that the PHD/Bromo domain of KAP1 associated with the central region of CTIP2 previously shown to mediate Tat binding (11). It should be noted that Tat has an additional CTIP2 binding site located at its C-terminal domain, between residues 717 and 813 (11). Coimmunoprecipitation experiments carried out with nuclear extracts expressing CTIP2, KAP1 and increasing amount of Tat, showed that Tat expression displaced CTIP2 from KAP1 suggesting that Tat may have a better affinity for KAP1 than CTIP2, (Figure 10 F, column 9 and 10).

**Figure 10:**
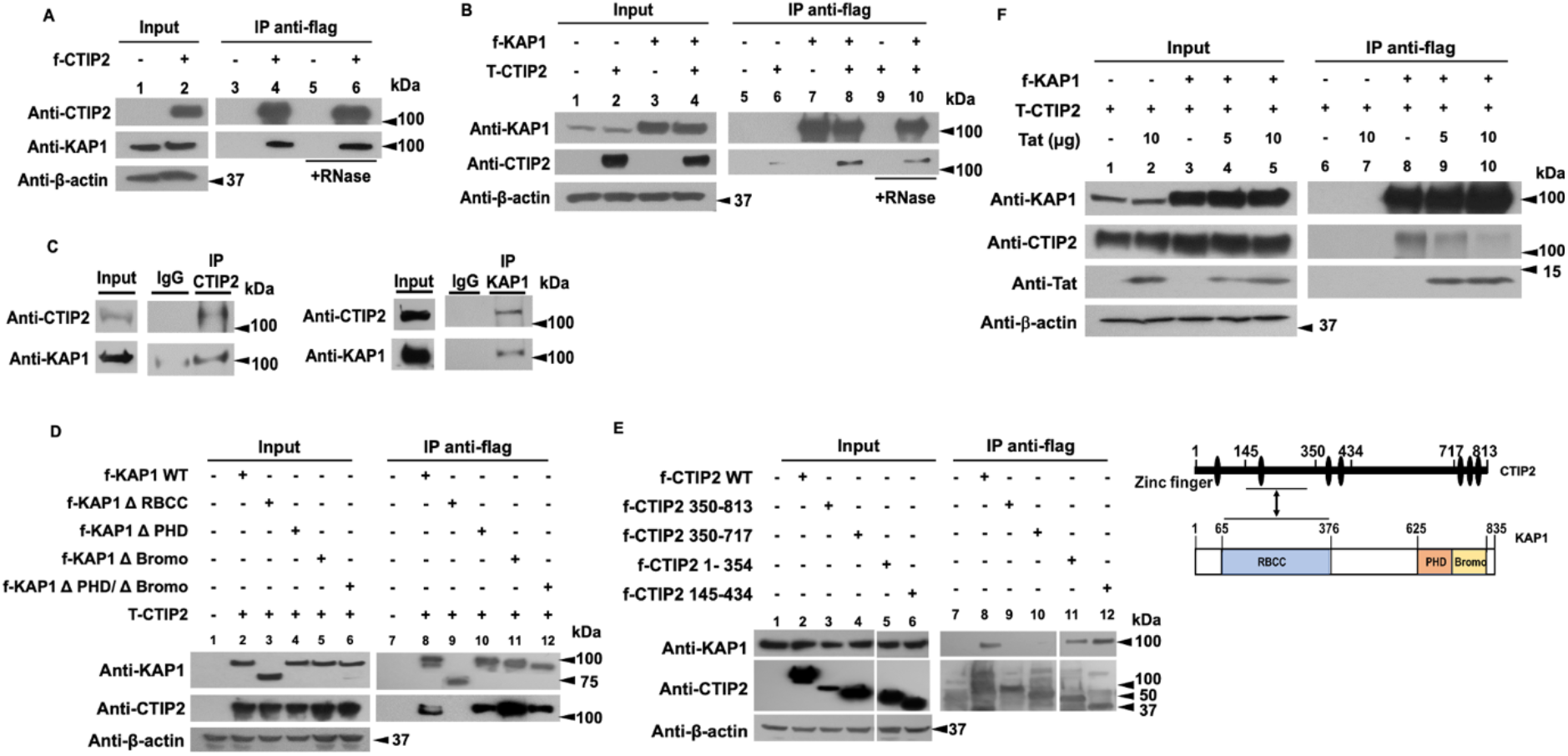
CTIP2 interacts with KAP1 in an RNA independent manner and competes with Tat for KAP1 binding. (A, B) HEK cells were transfected with flag-CTIP2 (A) or flag-KAP1 and Tap-CTIP2 (B). 48 h post-transfection, the nuclear protein extracts, treated or not with RNase, were subjected to immunoprecipitation with the flag antibody. The immunoprecipitated complexes were tested by Western blot for the presence of KAP1 (A) and CTIP2 (B). (C) The nuclear protein extracts of microglial cells were analyzed by Western blot for the expression of CTIP2 and KAP1 and submitted to immunoprecipitation experiments with CTIP2 and KAP1 antibodies. (D) The HEK cells were transfected with Tap-CTIP2 and one of the vectors encoding f-KAP1 WT: wild type, or KAP1 deleted from the RBCC domain (ΔRBCC), or PHD (ΔPHD), or Bromo (ΔBromo) or both PHD and Bromo domains (ΔPHD / ΔBromo), or with f-CTIP2 deletion mutants, as indicated (E). After 48h, the nuclear protein extracts were immunoprecipitated with flag antibody. The eluted protein complexes were analyzed by Western blot for the presence of CTIP2 and KAP1. (F) HEK cells were transfected with the indicated vectors in the presence of an increasing dose of Tat. After 48h, the nuclear protein extracts were treated as in D. The results are representative of at least two independent experiments.

## DISCUSSION

Antiretroviral therapy has significantly improved the management of HIV-1 infection. However, despite ART efficiency, it is still impossible to cure HIV-1 infected patients. Latently infected reservoirs are the major hurdles toward an HIV cure. Among the major reservoirs are the resting CD4+ T cells and monocyte-macrophage lineage, including microglial cells. These reservoirs escape immune surveillance and cannot be targeted by current therapies. HIV-1 gene transcription is silenced by epigenetic modifications. These epigenetic changes remain a major obstacle to the virus eradication and patients cure. We have previously shown that CTIP2 is a major player in the establishment and persistence of HIV-1 latency in microglial cells (9, 10, 12). In the present work we report the role and the mechanism of action of KAP1 as new partner of CTIP2 in the control of HIV-1 expression in myeloid cells. Functionally, KAP1 suppressed HIV-1 expression and contributed to viral latency in myeloid cells. In accordance, we found KAP1 associated with the latent HIV-1 promoter but not after reactivation. Interestingly, in CD4+ T cells, KAP1 binding to the HIV-1 promoter has been reported to be unsensitive to reactivations (31). These controversies suggest that the mechanisms underlying KAP1 functions may be cell type specific. It is now well established that the repressive form of KAP1 is SUMOylated on its Bromo domain (45) and associates with the NuRD and SETDB1 repressor complexes (reviewed in (46)). The repressors CTIP2 and KAP1 have multiple similarities. They both interact with the NuRD complex (47, 48), the DNA repair enzymes PARP1 and PRKDC (49, 50) and a repressive form of the P-TEFb complex including the 7SKsnRNP (26). CTIP2 and KAP1 have also common target genes like p21 (32, 51). Our results demonstrate a functional cooperation between the two repressors to repress HIV-1 gene transcription. Moreover, KAP1 and CTIP2 interact physically. We report that the RBCC domain of KAP1 associates with the central domain of CTIP2. CTIP2 association with LSD1 and HMGA1 has been described to inhibit Tat-dependent viral gene transcription and to promote its relocation in heterochromatin structures (10, 11, 13). Our results show that KAP1 cooperates with CTIP2 to repress Tat function. In addition, we found that KAP1 interacts and colocalizes with Tat in the nucleoplasm of microglial cells. The HIV-1 Tat transactivator is essential to the elongation step of the viral gene mainly because of its ability to recruit P-TEFb at the HIV-1 initiated transcripts. Thereby, inhibiting Tat activity results in hindering the positive feedback loop required for efficient HIV-1 transcription. Interestingly, Tat has been shown to be sensitive to proteasome degradation (39–44, 52). We observed that Tat expression was inversely proportional to KAP1 expression level suggesting that KAP1-mediated repression of Tat function results from Tat degradation. KAP1 has been shown to promote proteasomal degradation of targets proteins by its RBCC domain (53–55). Surprisingly, we observed that Tat degradation was not sensitive to RBCC depletion but mediated by the PHD and the Bromo domains of KAP1. In line with these results, the point mutation of the KAP1 Serine 824 located in the Bromo domain abrogated Tat degradation. The Bromo domain is known to be modified by SUMOylation giving to KAP1 its repressive form. The phosphorylation of Serine 824 causes a loss of this SUMOylation state and a loss KAP1 repressive activity (32). Thus, our results suggest that Tat degradation is promoted by repressive SUMOylated forms of KAP1. Finally, we show that KAP1 interacts with Tat with higher affinity than with CTIP2. Indeed, upon increasing Tat expression, CTIP2 is displaced from KAP1 associated complex suggesting that the dynamic of KAP1 association with CTIP2 depends on the virological status of the cells. Altogether, our results suggest that KAP1 is a major player in the control of HIV-1 gene expression in microglial cells, the main HIV-1 reservoir in the brain. Moreover, they further highlight the cell type specific mechanism of action of the repressors involved in viral latency. CTIP2 and KAP1 are highly expressed in resting T cells and depleted upon activation (31, 56). In addition, increased levels of CTIP2 are associated with persistence of HIV-1 latency in the brain (14) and increased level of KAP1 in the peripheral blood of gastric cancer patients is a biomarker predicting cancer stage progression (57–60). By transposition to what has been observed in oncology and virology, these repressors and their associated enzymatic activities may constitute good targets in therapies aiming at curing the HIV-1 infected patients

## ACKNOWLEDGEMENT

We thank Dr. David K. Ann, Dr. Ivan D’orso and Dr. Florence Cammas for providing vectors.

The authors wish it to be known that, in their opinion, the last three authors should be regarded as joint senior authors.

